# Relapse after intermittent access to cocaine: Discriminative cues more effectively trigger drug seeking than do conditioned cues

**DOI:** 10.1101/2024.02.22.580659

**Authors:** Ndeye Aissatou Ndiaye, Sema Abu Shamleh, Domiziana Casale, Sol’Abraham Castaneda-Ouellet, Isabel Laplante, Mike J. F. Robinson, Anne-Noël Samaha

## Abstract

When people with drug addiction encounter cues associated with drug use, this can trigger cravings and relapse. These cues can include conditioned stimuli (CSs) signaling drug delivery and discriminative stimuli (DSs) signaling drug availability. Compared to CS effects, DS effects are less explored in preclinical studies on cue-induced relapse. We compared CS and DS effects on reward seeking following abstinence from intermittent-access cocaine (or sucrose) self-administration. During 15-20 intermittent-access sessions, rats self-administered sucrose pellets or i.v. cocaine paired with a light-tone CS. Cocaine/sucrose was available for 5-min (signalled by a light; DS+) and unavailable for 25 min (signalled by different lighting conditions; DS-), and this cycled for 4 h/session. Following abstinence, we measured cocaine/sucrose seeking under extinction triggered by CS and DS presentation, and instrumental responding reinforced by these cues. Across intermittent-access sessions, rats increased lever pressing for cocaine or sucrose during DS+ periods and decreased responding during DS-periods. On days 2 and 21 of abstinence, only presentation of the DS+ or DS+ and CS combined elicited increased cocaine/sucrose-seeking behaviour (i.e., increased active lever presses). Additionally, presenting the DS-alongside the DS+ suppressed the increased cocaine-seeking behaviour otherwise produced by the DS+. Finally, on day 21 of abstinence, rats showed equivalent levels of lever pressing reinforced by the DS+, CS and by the DS+ and CS combined, suggesting comparable conditioned reinforcing value. After intermittent self-administration, cocaine-associated DSs and CSs acquire similar conditioned reinforcing properties, but DSs more effectively trigger increases in drug seeking.

## Introduction

The impact of cues on drug use and relapse is a critical focus in addiction treatment. Cocaine users commonly identify people, places and things associated with the drug amongst the most frequent triggers for craving and drug use (Shulman 1989). Laboratory studies confirm that drug-related cues lead to cravings and relapse in human drug users (O’Brien et al. 1992; Vafaie and Kober 2022) and stimulate drug-seeking behaviour in laboratory rats (Arroyo et al. 1999; de Wit and Stewart 1981; Panlilio et al. 1996; Shaham et al. 2003).

Animal models of cue-triggered drug seeking typically focus on relapse-like behaviour induced by conditioned stimuli, or CSs (Davis and Smith 1976; de Wit and Stewart 1981; Gueye et al. 2019; Meil and See 1996; 1997; Shaham et al. 2003; Shalev et al. 2002; Stewart et al. 1984). During the drug self-administration phase, rats associate a cue (e.g., a light/tone; the CS) with drug infusions, and after a period of abstinence, relapse to drug seeking is assessed by measuring the ability of that CS to reinforce instrumental responding in the absence of drug. Consequently, during both training and testing, the CS is contingent on the drug-seeking response. That is, the CS and the neural substrates it recruits operate *after* drug seeking is initiated. However, relapse also involves discriminative stimuli (DSs) that precede (i.e., trigger) drug-seeking actions, and that signal drug availability. DSs are very effective craving and relapse triggers, even months after the last bout of drug self-administration (Ciccocioppo et al. 2004; Madangopal et al. 2019; Weiss et al. 2001). Despite controlling drug seeking before and during relapse (Di Ciano and Everitt 2003), how DSs operate is largely unknown.

To better mimic human cocaine-taking patterns, which involve intermittency both between and within bouts of intake (Allain et al. 2015; Beveridge et al. 2012; Cohen 1994; Leri et al. 2004), many rat studies now use intermittent-access self-administration procedures (IntA). IntA involves alternating periods of cocaine availability (e.g., 5 min) and unavailability [e.g., 25 min; (Zimmer et al. 2011; Zimmer et al. 2012)]. Compared to continuous access procedures (e.g., Long or Short Access to cocaine), IntA is more effective in producing addiction-relevant features, despite lower total cocaine intake (Allain et al. 2017a; Allain et al. 2015; James et al. 2019; Kawa et al. 2019a; Samaha et al. 2021; Singer et al. 2018). This includes robust CS-induced cocaine-seeking behaviour (Gueye et al. 2019; James et al. 2019; Kawa et al. 2016; Kawa et al. 2019b; Nicolas et al. 2019; Singer et al. 2018), making IntA cocaine self-administration uniquely suited for studying the influence of cocaine-associated cues on relapse.

Two published studies have examined DS-induced relapse after IntA cocaine self-administration (Collins et al. 2022; Pitchers et al. 2017), with one other study also examining this with fentanyl (Martin et al. 2022). Pitchers et al. (2017) found that a cocaine-associated DS triggered significant cocaine seeking after intermittent cocaine use, especially in rats with a goal-tracking phenotype. However, they used extinction training before relapse testing, which might not reflect real-word conditions, where abstinence is forced (e.g., inpatient treatment) or voluntary. Relapse procedures without extinction might better capture the clinical situation (Epstein and Preston 2003; Fuchs et al. 2008), and they may also better capture the mechanisms of relapse, as different neural substrates mediate cue-induced cocaine seeking after extinction vs. after abstinence without extinction training (Augur et al. 2016; Fuchs et al. 2006; Muller Ewald et al. 2019). In the second IntA study, Collins et al. (2022) found that after intermittent cocaine use, rats endured electrical foot shock to seek cocaine when presented with a drug-associated DS (but not when a cocaine-paired CS was presented), with no sex differences in this effect. Thus, across the sexes, a cocaine-predictive DS can trigger relapse after IntA experience, even in the face of punishment.

Our study expands on this research by comparing DS vs. CS effects on cocaine or sucrose seeking behaviour after abstinence from IntA self-administration. We also assessed the conditioned reinforcing properties of these cues and examined the ability of a DS signaling cocaine unavailability to inhibit cocaine seeking.

## Materials and Methods

### Animals

All experimental procedures followed the guidelines of the Canadian Council of Animal Care and were approved by the animal care committee of the Université de Montréal. For Experiment 1, we used 16 male (7-8 weeks old; 200-225 g) and 16 female (7-8 weeks old; 150-175 g) Sprague Dawley rats (Envigo, Frederick barrier 208A, Maryland, USA). For Experiment 2, we used 7 male Sprague Dawley rats (21-22 weeks old, 500-600 g; Charles River, Barrier R04, Raleigh, NC). For Experiment 3, we used 40 female Sprague Dawley rats (7-8 weeks old; 150-175 g; Envigo, Frederick barrier 208A, Maryland, USA). All rats were housed 1/cage under a reverse dark/light cycle (12/12h with lights off at 8h30 AM). All self-administration training and testing occurred during the dark phase of the rat’s circadian cycle. Water was available *ad libitum*. During the 4 days of acclimatation after arrival to our institution, rats had free access to food (Charles River Purina chow #5075). From the 5^th^ day on, food was restricted to 18-20 g/day for females and 23-25 g/day for males (Algallal et al. 2020), except during i) sucrose self-administration training, where we gave 10 g/day to females and 15 g/day to males, and ii) the days surrounding catheter implantation surgery, where we gave 30 g/day to females and 33 g/day to males. Mild food restriction as used here promotes health in laboratory rats (Martin et al. 2010) and reduces the amount of cocaine used during drug self-administration sessions.

### Apparatus

All training and testing took place in standard operant cages (Med Associates, St Albans, VT) housed in sound-attenuating cabinets each with a ventilating fan. The front wall of each cage contained two retractable levers with a white cue light located above each and a recessed food/liquid port located between the levers. The back wall contained a white house light, and a Sonalert tone generator. On each training/testing day, rats were put in the operant cages and the house light and fan turned on and stayed on throughout each session. Two min later, both levers were inserted to signal the beginning of the session, and levers remained present throughout each session. Cages were cleaned between rats.

### Sucrose self-administration training

All rats were trained to press the active lever for 45-mg banana-flavoured sucrose pellets (product # 76285-260, VWR) during daily, 30-min sessions. For sessions 1-2, sucrose was available on a Fixed ratio 1 schedule of reinforcement (FR1). Rats had to earn a minimum of 20 pellets/session, on two consecutive sessions to proceed to the next training step. After two FR1 sessions, rats that did not meet this acquisition criterion (17/79) received a third FR1 session. Rats that still did not meet criterion after this 3^rd^ session (3/79 rats) received an overnight session (from 7 p.m. to 8 a. m. the following day) during which only the active lever was present. After acquisition under FR1, sucrose pellets were now available under an FR3 schedule of reinforcement during daily, 30-min sessions. After 2-3 FR3 sessions, all rats met the acquisition criterion and rats set to self-administer cocaine (Exps. 1 & 3) were prepared for catheter implantation, while rats set to self-administer sucrose (Exp. 2) proceeded to intermittent access (IntA) self-administration sessions.

### Catheter implantation

After the last sucrose self-administration training session, homemade catheters were implanted into the right jugular vein of rats under isoflurane anesthesia (5% for induction and 2% for maintenance) as in Samaha et al. (2011). They received a perioperative intramuscular injection of penicillin (0.02 ml Derapen, 300 mg/ml; CDMV, Saint-Hyacinthe, QC, Canada) and a subcutaneous injection of the analgesic Carprofen Rimadyl (0.03 ml of a 50 mg/ml solution; CDMV, QC, Canada). Rats were then left in their home cage for recovery for 10-14 days. During the first three days of recovery, skin incisions were cleaned daily with chlorhexidine and if needed, flamazine. Catheters were flushed daily with 0.05 ml saline or 0.05 ml heparin [0.1ml of a 0.2 mg/ml heparin (Sigma-Aldrich, Oakville, ON)/saline solution mixed with 2 mg/ml of baytril (enroflaxin CDMV, St Hyacinthe, Quebec, Canada)] on alternate days.

### Cocaine self-administration training

Starting 10-14 days after catheter implantation, rats in Exps. 1 and 3 were trained to self-administer intravenous cocaine hydrochloride infusions (0.5 mg/kg/infusion, delivered over 5 s) under an FR3 schedule of reinforcement, in 1-h daily sessions. Cocaine (Galenova, St-Hyacinthe, Quebec, Canada) was dissolved in 0.9% saline and filtered with corning bottle-top filters (0.22 μm PES membrane; Fisher Scientific, Whitby, ON, Canada). Every 3 days, the concentration of the cocaine solution was adjusted according to average rat weight. Each session started with activation of the cage house light and fan. Two min later, levers were inserted into the cage. Pressing the active lever produced a cocaine infusion without any experimenter-programmed drug-associated cues and no time out period. Acquisition criteria included taking a minimum of 6 infusions/session and making at least twice more active *vs*. inactive lever presses, on two consecutive days. Rats that met criteria (26/32 rats in Exp. 1, 33/40 in Exp. 3) began IntA self-administration sessions.

### IntA cocaine or sucrose self-administration with discriminative and conditioned stimuli

After self-administration training, rats received daily 4-h IntA sessions during which cocaine/sucrose was available during 5-min ON periods and then unavailable during 25-min OFF periods (Algallal et al. 2020; Allain et al. 2017a; Allain et al. 2021; Ndiaye et al. 2022; Zimmer et al. 2011; Zimmer et al. 2012). The cage house light and fan turned on at the beginning of each session. Two min later, the house light was turned off, and the levers were inserted into the cage. The light above the inactive lever was also illuminated to signal the first 5-min cocaine/sucrose ON period. This light thus served as a positive discriminative stimulus, or DS+. During the 5-min ON periods, pressing the active lever produced a sucrose pellet or cocaine infusion (0.5 mg/kg/inf, over 5 s) under FR3. Each earned reinforcer was also paired with a 5-s conditioned stimulus (CS) consisting of illumination of the cue light above the active lever and sounding of a tone (2900 Hz, 75 dB). After each 5-min cocaine/sucrose ON period, the cue light extinguished, signaling a 25-min OFF period, during which lever pressing had no consequence. Thus, extinction of lighting in the chamber served as a negative discriminative stimulus, or DS-(Exps. 1 & 2). Previous work shows that illumination and extinction of discrete lights serve as effective discriminative stimuli for IntA cocaine self-administration (Collins et al. 2022). As detailed below under ‘Experiment 3: Conditioned inhibition of cocaine seeking’, in Exp. 3, the DS-consisted of illumination of the light above the active lever, rather than extinction of all ambient lighting. During each IntA session, the sequence of 5-min ON and 25-min OFF periods was repeated 8 times, resulting in a 4-h session. In each experiment, rats received the number of IntA sessions needed for average responding for cocaine/sucrose to peak during ON/DS+ periods and to diminish during OFF/DS-periods. Thus, rats in Exps. 1 & 3 received 14-15 IntA sessions and rats in Exp. 2 received 10.

### Catheter patency

In Exps. 1 & 3, we verified catheter patency at the end of the last IntA session by administering propofol (0.05 ml of a 10 mg/ml solution, i.v.; Fresenius Kabi Canada Ltd, Richmond Hill, ON) to each rat, followed by 0.05 ml of saline. Rats that did not become ataxic after 1 dose of Propofol were immediately given a 2^nd^ dose. Those that did not become ataxic in response to the 2^nd^ dose were excluded from statistical analyses.

### Cue-induced cocaine/sucrose seeking

On days 2 and 21 days following the last IntA session, we assessed the effects of the DS+, DS-, CS, DS+CS combined (the latter combination in Exps. 1 & 2 only) and DS+DS-combined (Exp. 3) on cocaine/sucrose seeking. The cues were presented independent of the rat’s behaviour, to measure the ability of these cues to trigger increases in reward seeking. All rats were tested at both abstinence timepoints, and rats stayed in their home cages between tests. Each test began with a 10-min period (2-min period in Exp. 3) during which the house light and fan were turned on, and levers were present, but no cocaine/sucrose cues were presented. After this baseline period, the house light was extinguished and we gave alternating, response-independent presentations of the cues in pseudo randomized order. Cue presentations lasted 2 min each, with a 2-min inter-trial interval during which the house light was turned on. Each cue was presented 4 times in pseudo-random order and each test lasted 74 min.

## Experiment 1: Discriminative *vs* conditioned stimulus effects on cocaine seeking

### IntA cocaine self-administration with discriminative stimuli

Here we assessed the effects of cocaine-associated discriminative and conditioned stimuli on cocaine-seeking behaviour after a history of IntA cocaine self-administration (Figs. 1A-B). After sucrose self-administration training, male (n = 16) and female (n = 16) rats self-administered cocaine under an IntA schedule, as described above. On Session 14, average responding for cocaine followed a reliable pattern whereby lever pressing peaked during DS+ periods and decreased during DS-periods. This suggested that the DSs controlled cocaine self-administration behaviour. Alternatively, the rats may have been keeping time during each session. To address this, we gave rats 4 more IntA sessions that now included cocaine OFF/DS-periods of varying length. Thus, during IntA sessions 15-18, rats now experienced two 20-min, four 25-min and two 30-min OFF/DS-periods, in random order. If the DSs controlled self-administration behaviour, responding should still peak during DS+ periods and dip during DS-periods. If instead rats were keeping time, then this would be indicated by a ramping up of responding during DS-periods when these periods are longer than expected. During the IntA phase, 4/16 males and 2/16 females were excluded from further testing and data analysis either because they did not reliably self-administer cocaine or did not reliably discriminate between DS+/ON and DS-/OFF periods. In addition, 2/16 males and 4/16 females lost catheter patency during this period and were excluded from subsequent testing and data analysis.

**Figure 1:**
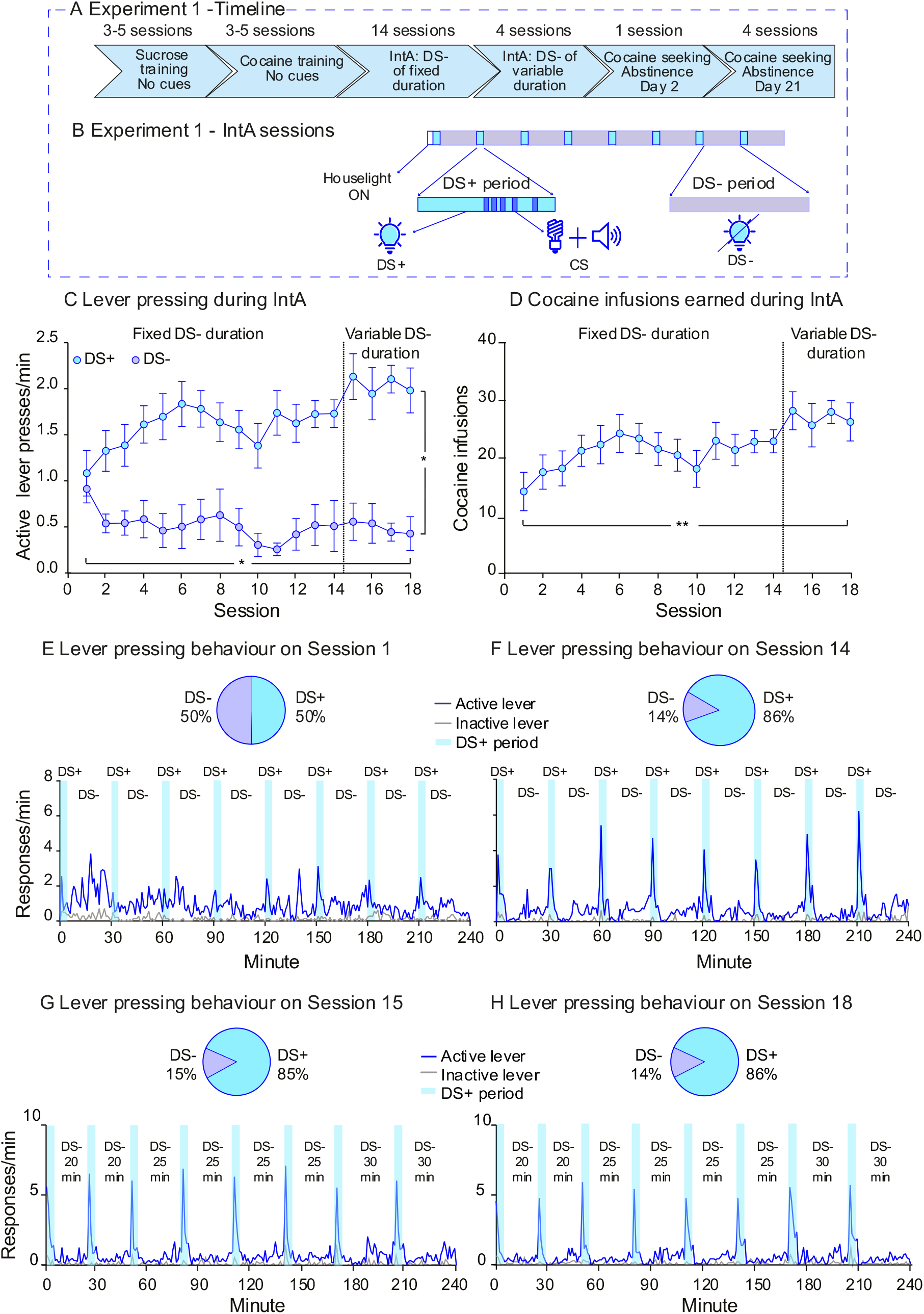
Intermittent access cocaine self-administration (IntA) under the control of discriminative stimuli signaling drug availability (DS+) vs. unavailability (DS-) in Experiment 1. **A** The sequence of experimental events. **B** IntA sessions alternated between 5-min cocaine ON periods and 25-min cocaine OFF periods. Cocaine available/ON periods were signaled by activation of the light above the inactive lever (DS+), and each cocaine infusion earned during these periods was paired with activation of the light above the active lever and sounding of a tone (conditioned stimulus, or CS). Cocaine unavailable/OFF periods were signaled by extinction of all lighting in the test cage (DS-). **C** Over sessions, rats learned to press more on the active lever during DS+ periods than during DS-periods. **D** Rats escalated their cocaine intake across sessions. **E** On IntA Session 1, rats did not show discriminative responding during DS+ versus DS-periods. **F** By IntA Session 14, rats pressed most on the active lever in response to the DS+ and decreased their responding during DS-presentation. **G, H** Responding on the active lever continued to peak during DS+ periods and dip during DS-periods even when DS-periods were of unpredictable length. Note that the 20-, 25- and 30-min DS-periods are presented in ascending order in **G-H** for simplicity, however they occurred randomly during Sessions 15-18. Data are mean ± SEM, n = 14.

### Cue-induced cocaine seeking

On days 2 and 21 days after the last IntA session, the remaining rats were tested for cocaine seeking induced by response-independent presentation of the DS+, DS-, CS and DS+CS combined (Fig. 2A), as described above.

**Figure 2:**
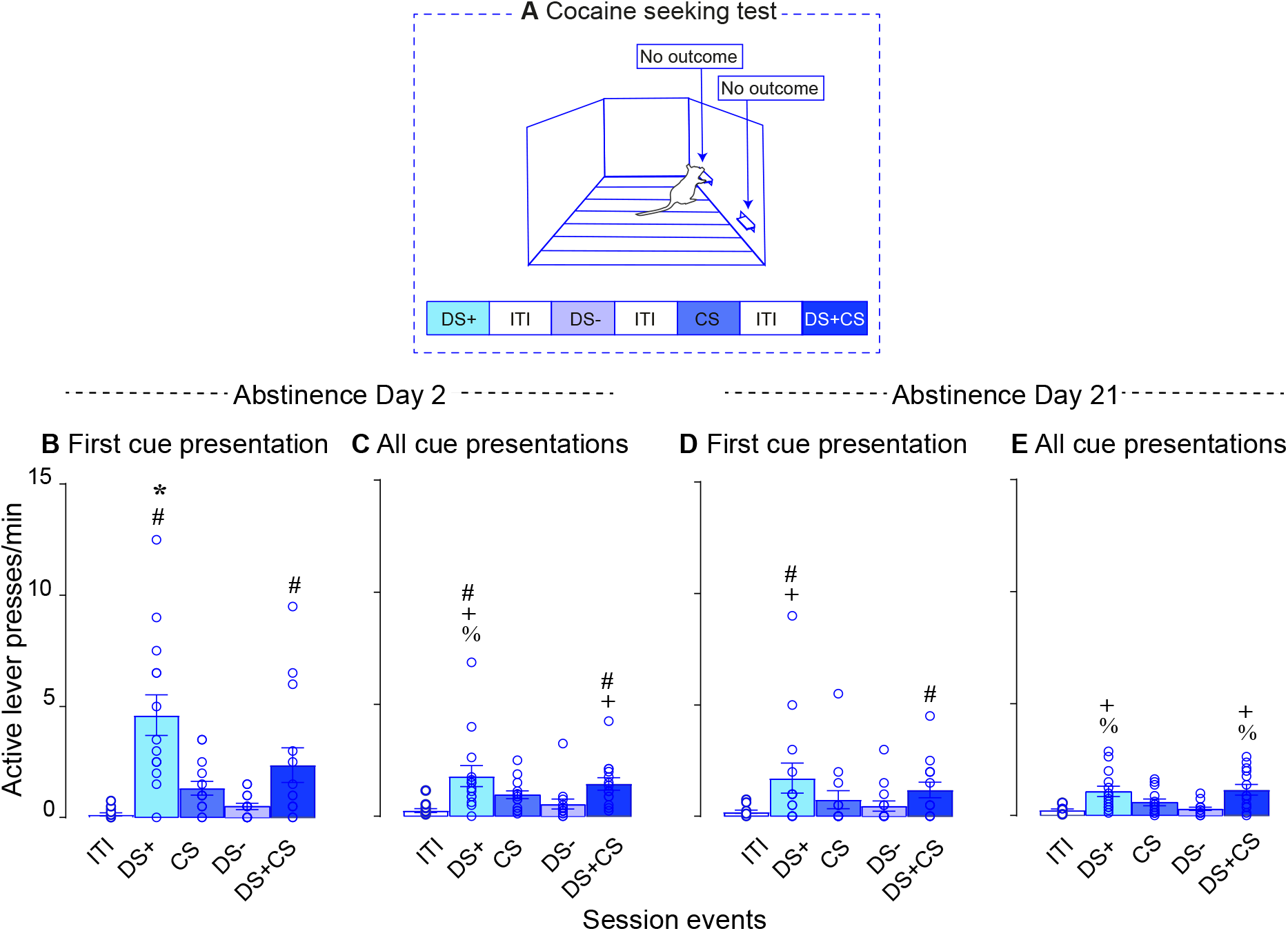
At both early (Day 2) and later abstinence times (Day 21), the cocaine-predictive discriminative stimulus (DS+) triggered the highest levels of cocaine-seeking behaviour. **A** During the cocaine-seeking tests, all cues (DS+, CS, DS- and DS+ and CS combined) were presented independent of responding and pressing on the levers produced no outcome (no cues and no cocaine). **B, C** On Day 2 of abstinence, DS+ presentation triggered higher rates of cocaine-seeking behaviour compared to the other cues. **D, E** On Day 21 of abstinence, only DS+ presentation—alone or alongside the CS—triggered significant cocaine-seeking behaviour. ITI; inter-trial interval; CS; conditioned stimulus. Data are mean ± SEM, n = 14; * vs. all other conditions; # vs. ITI; + vs. DS-; % vs. CS.

### Tests for conditioned reinforcement

Finally, to assess the conditioned reinforcing properties of the cocaine-associated cues, we gave rats test sessions during which they could lever press for presentations of the DS+, CS and DS+CS combined (1 cue type/session, counterbalanced; Fig. 3A). These tests were run on consecutive days, 22 days after the last IntA session. Each test started with a 2-min period during which the house light and fan were turned on and levers were inserted. After this 2-min period, the house light was extinguished and now pressing on the active lever produced a 5-s presentation of the DS+, CS or DS+CS, under FR1. These sessions lasted 20 min.

**Figure 3:**
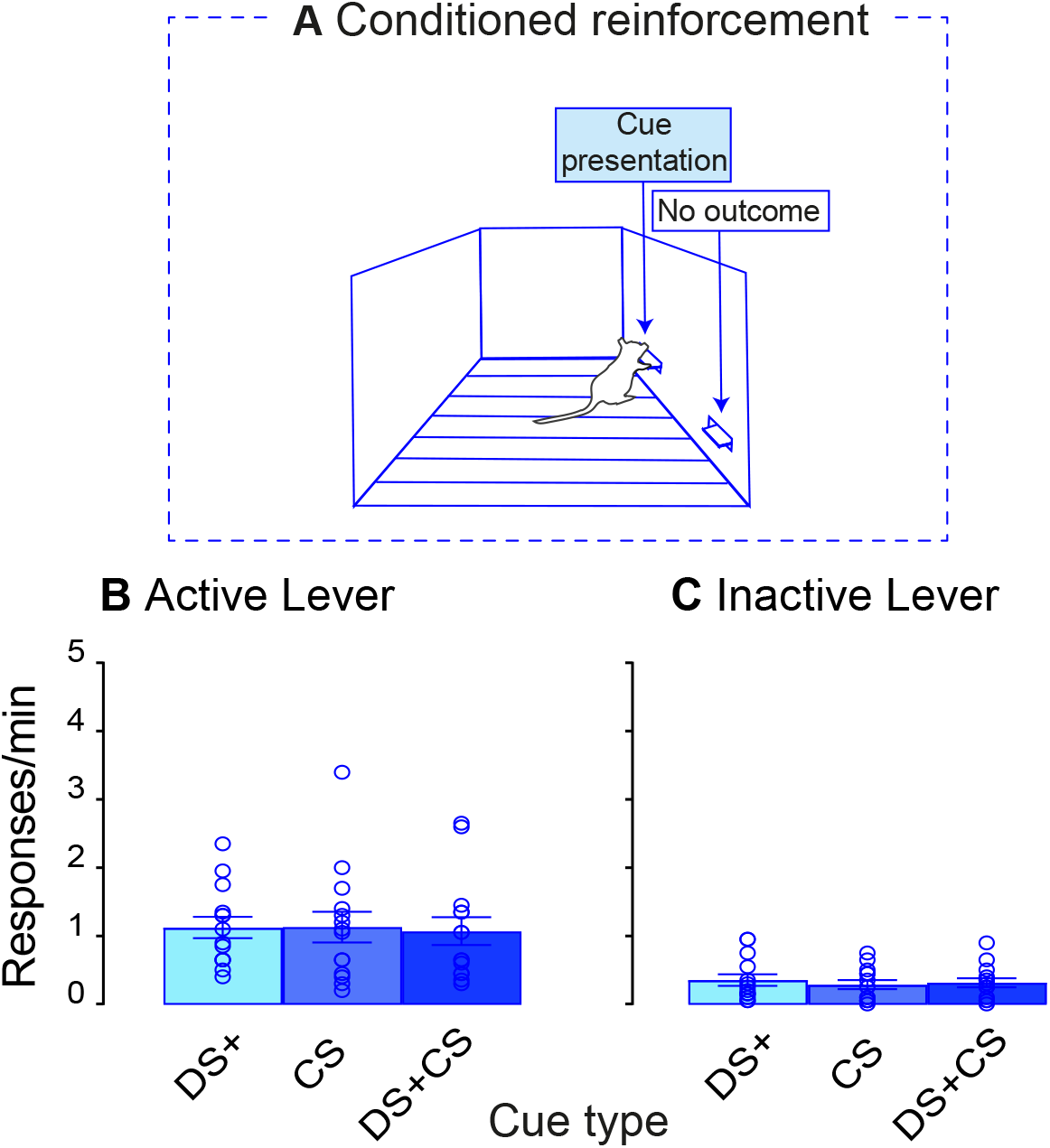
A cocaine discriminative stimulus (DS+) and a cocaine conditioned stimulus (CS), presented alone or together, have similar conditioned reinforcing effects. **A** Starting on abstinence Day 22, rats received test sessions during which pressing on the active lever produced a 5-s presentation of the DS+, CS or DS+CS combined (no cocaine was delivered; one cue type/session). **B, C** Rats pressed more on the active vs. inactive lever during conditioned reinforcement tests, and rats also responded similarly for the DS+, CS and DS+CS combined. Data are mean ± SEM, n = 14.

## Experiment 2: Discriminative *vs* conditioned stimulus effects on sucrose seeking

### IntA sucrose self-administration with discriminative stimuli

To determine whether effects observed in Exp. 1 generalize to a non-drug reinforcer, here we assessed the effects of discriminative and conditioned stimuli on sucrose-seeking behaviour after IntA sucrose self-administration (Figs. 4A-B). After sucrose self-administration training, male rats (n = 7) self-administered sucrose pellets during IntA sessions, as described above. Procedures were similar to those in Exp. 1, with the following exception. Rats received 10 initial IntA sessions of sucrose self-administration, because this was sufficient for average responding to reliably peak during DS+ periods and dip during DS-periods. Rats then received 10 more IntA sessions during which OFF/DS-periods were of variable length, as described for Exp. 1.

**Figure 4:**
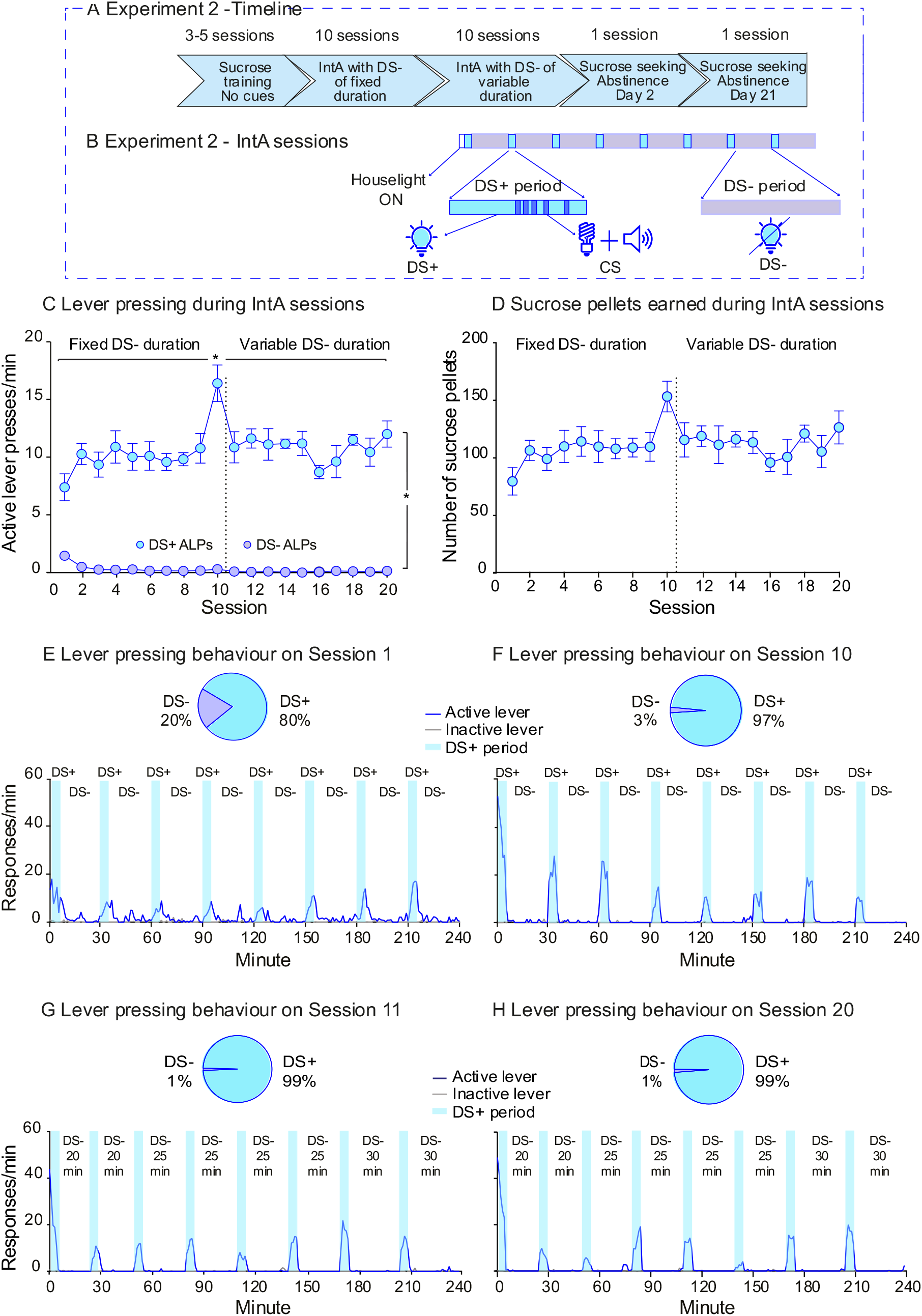
Intermittent access sucrose self-administration (IntA) under the control of discriminative stimuli signaling sucrose availability (DS+) vs. unavailability (DS-) in Experiment 2. **A** The sequence of experimental events. **B** IntA sessions alternated between 5-min sucrose ON periods and 25-min sucrose OFF periods. Sucrose ON periods were signaled by activation of the light above the inactive lever (DS+), and each earned sucrose delivery during these periods was paired with a conditioned stimulus (CS) consisting of activation of the light above the active lever and sounding of a tone. Sucrose OFF periods were signaled by extinction of all lighting in the cage (DS-). **C** Over sessions, rats learned to press more on the active lever during DS+ periods than during DS-periods. **D** Rats maintained a relatively stable level of sucrose intake across the 20 IntA sessions. **E** On IntA Session 1, rats already showed discriminative responding, pressing most on the active lever during DS+ periods. **F** By IntA Session 14, discriminative responding was further enhanced, with rats showing peaks in responding during the DS+ and virtually no responding during the DS-. **G, H** Responding on the active lever continued to pea during the DS+ and dip during the DS-, even when DS-periods were of unpredictable length. Note that the 20-, 25- and 30-min DS-periods are presented in ascending order in G-H for simplicity, however they occurred randomly during Sessions 11-20. Data are mean ± SEM, n = 7; in **C**, * *p* < 0.05 for DS effect and Session effect.

### Cue-induced sucrose seeking

On days 2 and 21 days after the last IntA session, we assessed sucrose-seeking behaviour induced by response-independent presentation of the DS+, DS-, CS and DS+CS combined (Fig. 5A), as described above.

**Figure 5:**
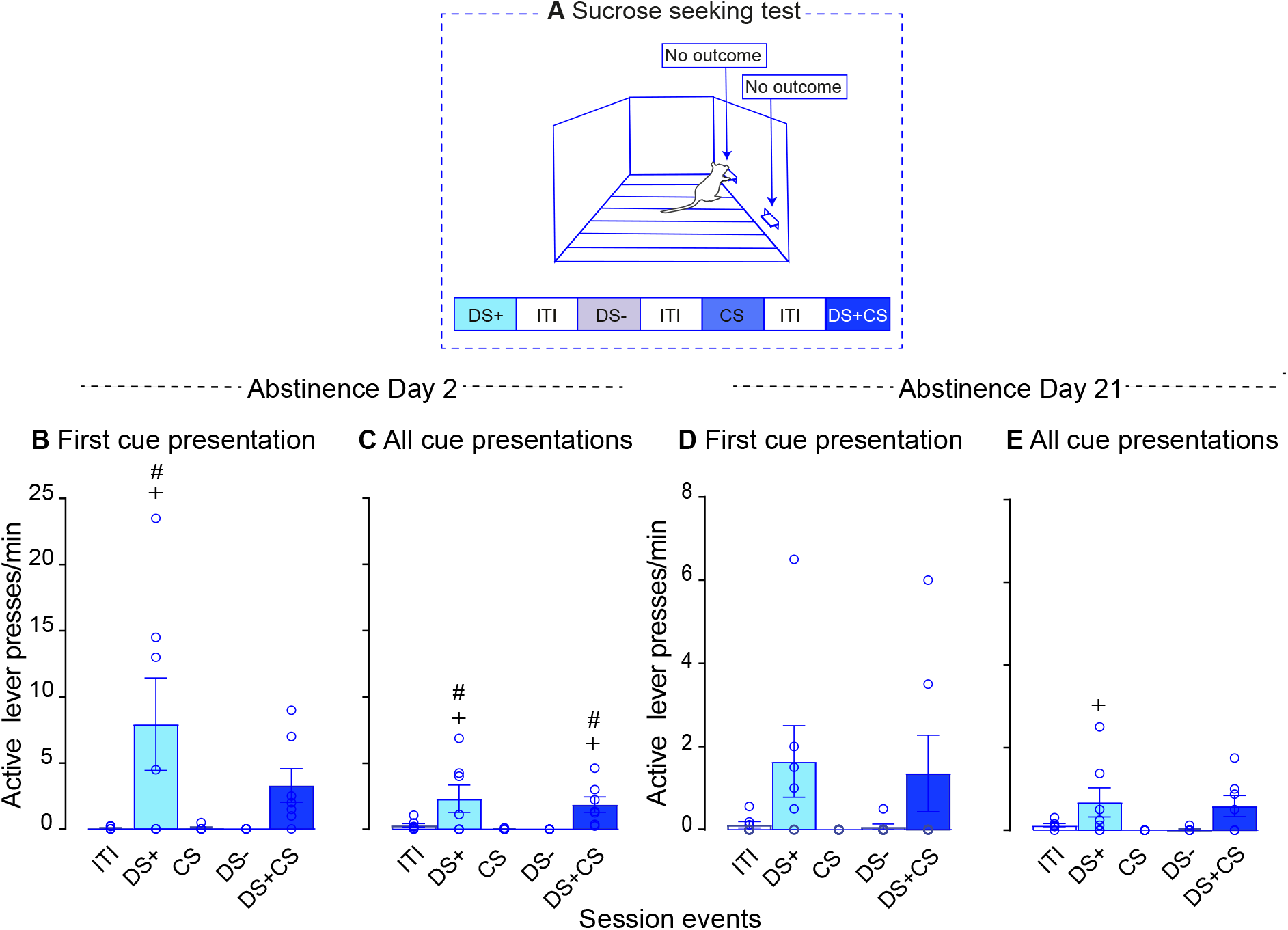
The sucrose-predictive discriminative stimulus (DS+) triggered the highest levels of sucrose-seeking behaviour after abstinence. **A** During the sucrose-seeking tests, all cues (DS+, CS, DS- and DS+ and CS combined) were presented independent of responding and pressing on the levers produced no outcome (no cues and no sucrose). **B, C** On abstinence Day 2, DS+ presentation triggered higher rates of cocaine-seeking behaviour compared to the other cues. **D, E** On abstinence Day 21, DS+ presentation triggered more sucrose-seeking behaviour compared to the CS and DS-. ITI; inter-trial interval; CS; conditioned stimulus. Data are mean ± SEM, n = 7; # vs. ITI; + vs. CS and DS-.

## Experiment 3: Conditioned inhibition of cocaine seeking

### IntA cocaine self-administration with discriminative stimuli

In Exp. 3 (Figs. 6A-B) we determined the extent to which a DS-acts as a conditioned inhibitor of cocaine relapse after IntA cocaine experience. An additional goal was to determine the effects of a DS+ and a DS-on responding for cocaine when the two stimuli are of the same modality (i.e., illumination of a discrete cute light in both cases). Procedures were as in Exp. 1 with the following exceptions: *i)* the DS-was the light above the active lever, and *ii)* the CS (5 s) was a tone (2900 Hz, 75 dB) and flashing house light (1 Hz). Forty female rats received 15 IntA sessions, all with 5-min cocaine ON/DS+ periods and 25-min cocaine OFF/DS-periods. During the IntA phase, 5 rats were excluded because they did not reliably self-administer cocaine or did not reliably discriminate between DS+/ON and DS-/OFF periods and 7 rats were excluded because they lost catheter patency.

**Figure 6:**
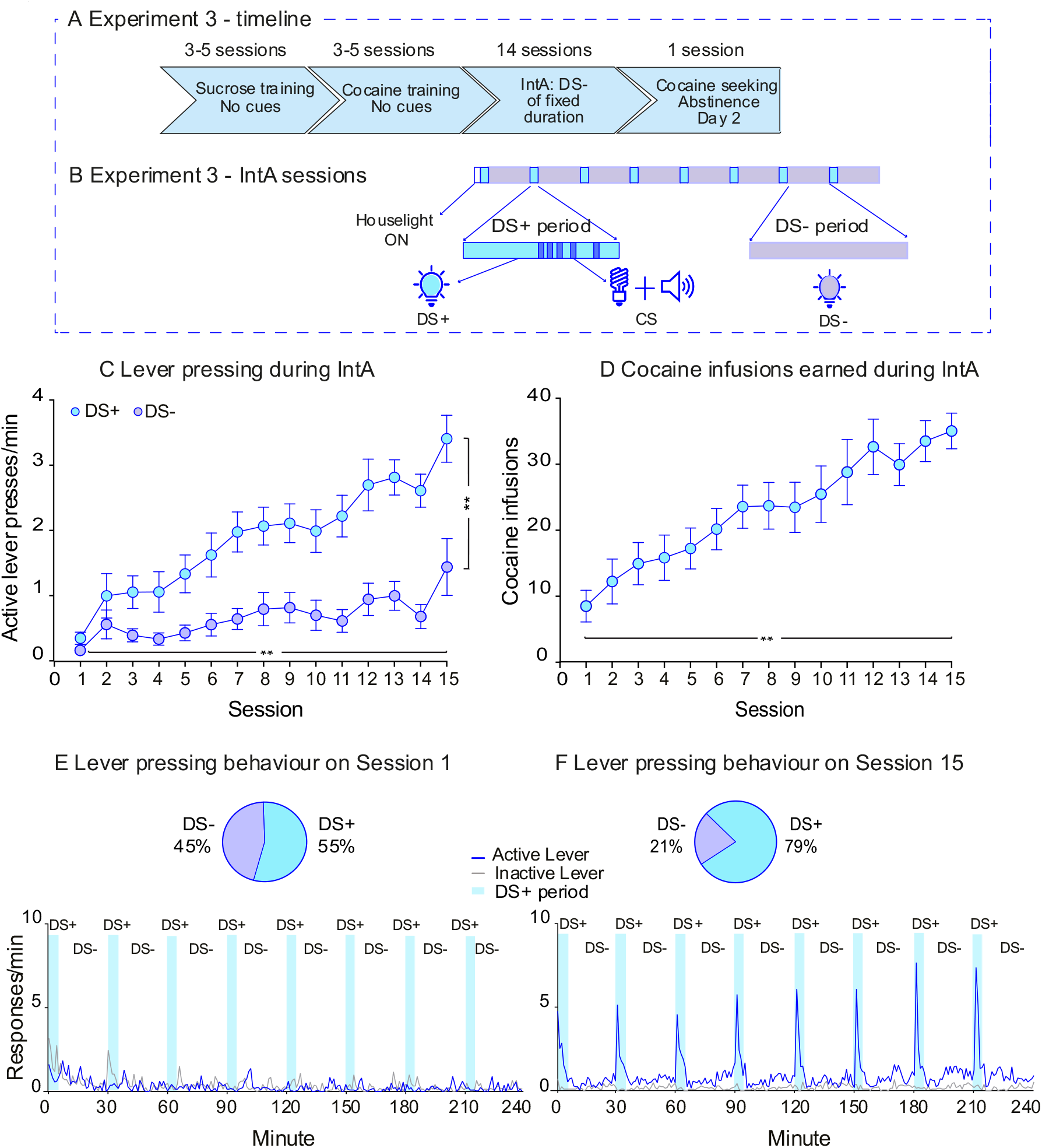
Discriminative stimuli signaling drug availability (DS+) vs. unavailability (DS+) control cocaine self-administration under intermittent access (IntA) conditions in Experiment 3. **A** The sequence of experimental events. **B** IntA sessions alternated between 5-min cocaine ON periods and 25-min cocaine OFF periods. Illumination of the light above the inactive lever (DS+) signaled cocaine ON periods, and during these periods, each earned cocaine infusion was paired with a conditioned stimulus (CS) consisting of activation of a flashing house light and sounding of a tone. Illumination of the light above the active lever (DS-) signaled cocaine OFF periods. **C** Over sessions, rats learned to press more on the active lever during the DS+ than during the DS-. **D** Rats escalated their cocaine intake across the 15 IntA sessions. **E** On IntA Session 1, rats responded on the active lever at comparable rates during DS+ and DS-periods. **F** By IntA Session 15, rats pressed most on the active lever during DS+ periods and inhibited their responding during DS+ periods. Data are mean ± SEM, n = 21. In **C**, ** *p* < 0.0001 for DS effect and Session effect; in **D** ** *p* < 0.0001 for Session effect.

### Cue-induced cocaine seeking

Two days after the 15^th^ session, rats received a cue-induced cocaine-seeking test with response-independent presentations of the DS+, DS-, CS and DS+DS-combined (Fig. 7A). If the DS-acts as a conditioned inhibitor of cocaine seeking, it should suppress the excitatory effects of a DS+ on responding for cocaine during the cocaine-seeking test.

**Figure 7:**
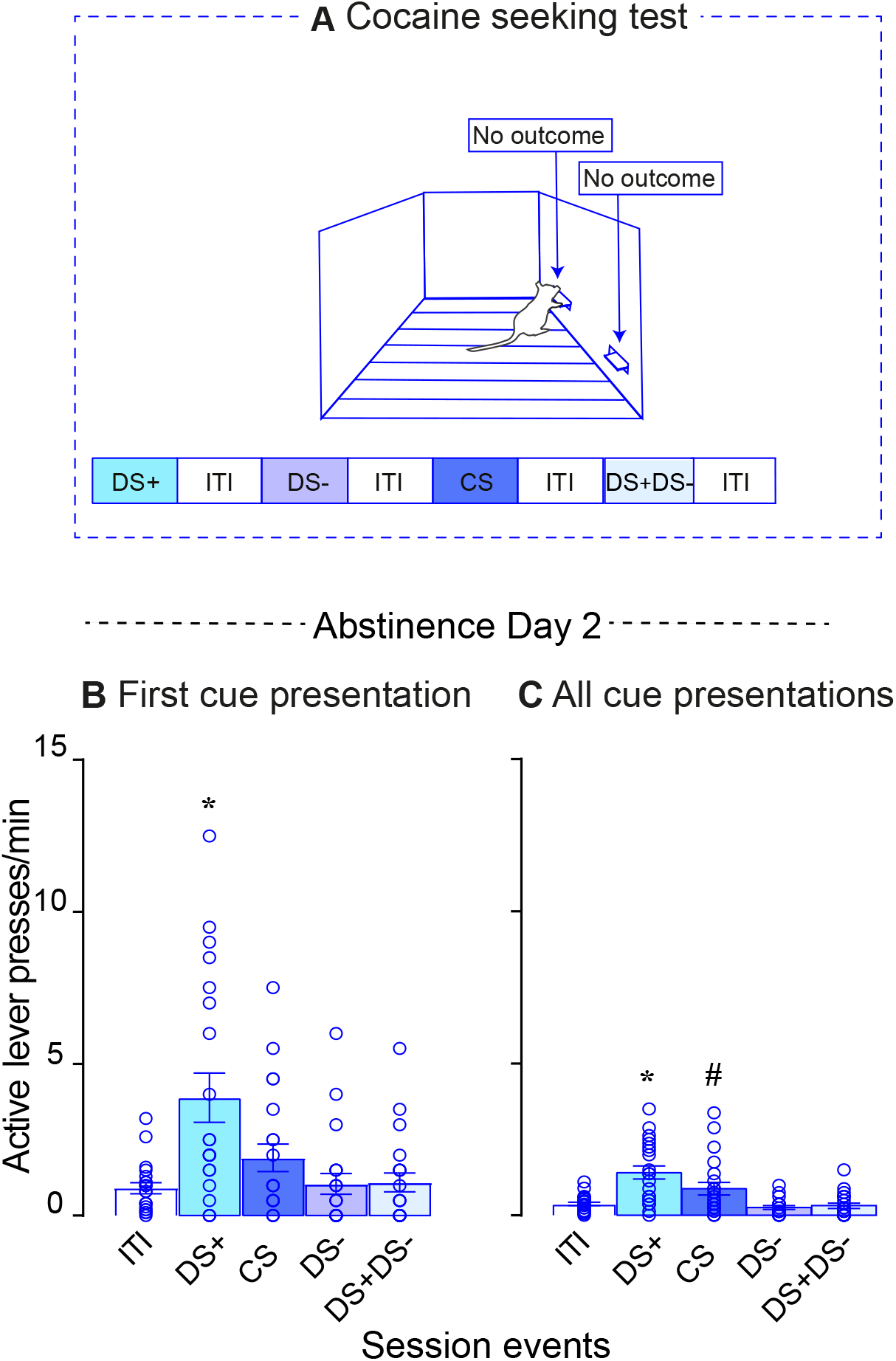
Conditioned inhibition of cocaine-seeking behaviour by a discriminative stimulus signaling cocaine unavailability (DS-). **A** On Day 2 of abstinence from cocaine, rats received a cocaine-seeking test during which all cues (DS+, CS, DS- and the DS+ and DS-combined) were presented independent of responding. Pressing on the levers produced no outcome (no cues and no cocaine). **B, C** During the cocaine-seeking test, DS+ presentation triggered higher rates of cocaine-seeking behaviour compared to the other cues, and this effect was inhibited by presenting the DS-alongside the DS+. ITI; inter-trial interval, CS; conditioned stimulus. Data are mean ± SEM, n = 21. * vs. all other conditions; # vs. ITI.

### Data analysis

Data were analysed with GraphPad Prism 8 and SPSS 20. The alpha level was set to *p* ≤ 0.05. During IntA sessions, we used two-way repeated-measures analysis of variance (ANOVA) to analyse the effects of the DSs on lever pressing/min (DS type x Session; both as within-subjects variables), we used one-way ANOVA to analyse cocaine intake across sessions and we used paired *t*-tests to assess escalation of intake. In Exp. 1, we used three-way mixed ANOVA to compare the sexes on active lever pressing during IntA sessions (Sex x DS type x Session; the latter two as within-subjects variables. For cocaine/sucrose seeking tests, we used two-way repeated measures ANOVA to analyse the effects of cue type on lever pressing/min (Cue type x Lever type). In Exp. 1, we used mixed three-way ANOVA to compare the sexes on lever pressing behaviour during the seeking tests (Sex x Cue type x Lever type; the latter two as within-subjects variables). When main or interaction effects were significant, these were analysed further using Holm-Sidak tests. Data in figures are expressed as mean values ± SEM.

## Results

### Experiment 1: Discriminative *vs* conditioned stimulus effects on cocaine seeking

#### IntA cocaine self-administration with discriminative stimuli

Active lever pressing during IntA sessions was similar across the sexes, and the two sexes also showed similar patterns of responding to the CS and DSs during cocaine-seeking tests and conditioned reinforcement tests (all *P*’s > 0.05). Thus, we pooled data across the sexes for analysis. Figs. 1C-D show cocaine self-administration behaviour across the 18 IntA sessions. During sessions 1-14, DS-periods were of fixed duration (i.e., 25 min), and during sessions 15-18 DS-periods lasted 20, 25 or 30 min. Rats pressed more on the active lever during cocaine ON/DS+ vs. cocaine OFF/DS-periods and this discrimination increased across sessions (Fig. 1C; DS type; *F* (1, 13) = 82.80, *p* < 0.0001; Session x DS type; *F* (17, 221) = 3.60, *p* < 0.0001). Cocaine intake also escalated across the 18 sessions (Fig. 1D; Session effect; F (17, 221) = 2.58, *p* = 0.0008; Session 1 vs. Session 18; *t* (13) = 2.35, *p =* 0.02), increasing by 381% (SEM ± 142) from Session 1 to 18. Thus, rats learned that the DS+ signaled cocaine availability, and the DS-signaled cocaine unavailability.

Figures 1E-H show the progression of discrimination training across sessions by illustrating the rate of responding on the levers. On Session 1, the rats pressed more on the active vs. inactive lever (Fig. 1E; Lever type; *F* (1, 13) = 27.81, *p* = 0.0002), and the rate of active lever pressing varied across the session (Time effect; *F* (240, 3120) = 1.19, *p* = 0.02). During this first IntA session, active lever presses were evenly distributed across DS+ and DS-periods (pie chart inset in Fig. 1E; 50 % for DS+ and 50 % for DS-; *t* (13) = 0.06, p = 0.48; SEM ± 3.89).

Similarly, on Session 14, rats still pressed most on the active lever (Fig. 1F; Lever type; *F* (1, 13) = 7.87, *p* = 0.01) and the rate of active lever pressing also varied across the session (Fig. 1F; Time effect; *F* (240, 3120) = 4.5, *p* < 0. 0001). Importantly, on Session 14, rats made most of their active lever presses during DS+ periods and suppressed their responding during DS-periods (pie chart inset in Fig. 1F; 86 % for DS+ and 14 % for DS-, SEM ± 4.87; *t* (13) = 7.26, *p* < 0.0001).

The response peaks during DS+ periods could be because the DSs controlled cocaine self-administration behaviour or alternatively, because the rats were keeping time during each session. To address this, we varied the duration of the DS-periods for 4 IntA sessions. On the first such session (Session 15, Fig. 1G), responding remained low during DS-periods and peaked during DS+ periods and the majority of responding occurred during DS+ periods (Lever type; *F* (1, 13) = 14.53, *p* = 0.002; Time effect, *F* (240, 3120) = 6.5, *p* < 0.0001; pie chart inset in Fig. 1G; 85 % for DS+ and 15 % for DS-, SEM ± 3.45; *t* (13) = 10.24, *p* < 0.0001). This pattern of responding was maintained through Session 18 (Fig. 1H; Lever type; *F* (1, 13) = 15.76, *p* = 0.002; Time effect, *F* (240, 3120) = 4.67, *p* < 0.0001; pie chart inset; 86 % for DS+ and 14 % for DS-, SEM ± 4.38; *t* (13) = 8.12, *p* < 0.0001). Thus, during IntA sessions rats were not pacing their cocaine self-administration behaviour using an internal timer, and instead responding was under the control of the discriminative stimuli signaling drug availability vs. unavailability.

### Cue-induced cocaine seeking

On Days 2 and 21 following the last IntA session (session 18), we assessed cocaine-seeking behaviour induced by response-independent presentation of the DS+, DS-, CS and DS+CS combined. On each test session, each cue was presented 4 times, and lever pressing produced no outcome (Fig. 2A; no cocaine or cues). We analysed the rate of responding to the first presentation of each cue (Figs. 2B and D) and across the 4 presentations of each cue (Figs. 2C and E). On Day 2 of forced abstinence, rats responded most on the previously cocaine-associated (active) lever, and this effect was dependent on cue type, both when we analysed responding to the first presentation of each cue (Fig. 2B shows active lever presses and Supplemental Fig. 1A shows inactive lever presses; Lever type, *F* (1, 13) = 25.95, *p* = 0.0002; Cue type *F* (4, 52) = 9.71, *p* < 0.0001; Lever x Cue *F* (4, 52) = 10.35, *p* < 0.0001) and responding across the 4 presentations of each cue (Fig. 2C shows active lever presses and Supplemental Fig. 1B shows inactive lever presses; Lever type, *F* (1, 13) = 33.67, *p* < 0.0001; Cue type *F* (4, 52) = 8.5, *p* < 0.0001; Lever x Cue *F* (4, 52) = 8.21, *p* < 0.0001). Cue type did not influence inactive lever presses (Supplemental Figs. 1A-B; all *p* values > 0.05). However, cue type determined responding on the previously cocaine-associated lever. First, only presentations of the DS+ and DS+CS combined increased responding on the previously cocaine-associated lever relative to the inter-trial interval (Fig. 2B; DS+ v. ITI, *p* < 0.0001; DS+CS v. ITI, *p* = 0.002; Fig 2C; DS+ v. ITI, *p* < 0.0001; DS+CS v. ITI, *p* = 0.001). Second, DS+ presentation produced higher rates of responding compared to all other cue types (Fig. 2B; DS+ v. DS-, *p* < 0.0001; DS+ v. CS, *p* < 0.0001; DS+ v. DS+CS, *p* = 0.002; DS+CS v. DS-, *p* = 0.01; Fig 2C; DS+ v. DS-, *p* < 0.0001; DS+ v. CS, *p* = 0.0003; DS+CS v. DS-, *p* = 0.0003).

After 21 days of forced abstinence, responding was highest on the previously cocaine-associated lever, and responding also varied as a function of cue type (Fig. 2D shows active lever presses and Supplemental Fig. 1C shows inactive lever presses; Lever type, *F* (1, 13) = 11.12, *p* = 0.005; Cue type *F* (4, 52) = 1.68, *p* = 0.17; Lever x Cue, *F* (4, 52) = 3.6, *p* = 0.01; Fig. 2E shows active lever presses and Supplemental Fig. 1D shows inactive lever presses; Lever type, F (1, 13) = 6.02, *p* = 0.03; Cue effect, F (4, 52) = 4.95, *p* = 0.002; Lever x Cue effect *F* (4, 52) = 4.63, *p* = 0.003). Only DS+ and DS+CS presentation increased responding on the previously cocaine-associated lever relative to the inter-trial interval, and presentation of the DS+ and DS+CS also produced higher rates of responding compared to the other cue types (Fig. 2D; DS+ v. ITI, *p* = 0.001, DS+ v. DS-, *p* = 0.009; Fig. 2E; DS+ v. ITI, *p* = 0.0006; DS+ v. CS, *p* = 0.002; DS+ v. DS-, *p* < 0.0001; DS+CS v. ITI, *p* = 0.0002; DS+CS v. CS, *p* = 0.0008; DS+CS v. DS-, *p* < 0.0001). Cue type did not influence inactive lever presses (Supplemental Figs. 1C-D; all *p* values > 0.05).

### Conditioned reinforcement

Between days 22 and 24 of forced abstinence from cocaine, we determined the extent to which the DS+, CS and DS+CS combined acquired conditioned reinforcing value, by allowing rats to lever press to earn presentations of these cues, in the absence of cocaine (Fig. 3A; 1 test session/cue type). Across cue types, rats pressed more on the active vs. inactive lever (Figs 3B-C; Lever type, *F* (1, 13) = 36.06, *p* 0.0001). Responding did not depend on cue type (Figs. 3B-C; Cue type, *F* (2, 26) = 0.06, *p* = 0.94; Lever x Cue, *F* (2, 26) = 0.08, *p* = 0.92). Thus, rats were willing to work equally for presentations of a DS that previously signaled cocaine availability, a CS that previously signaled cocaine delivery, or the two cues combined. This suggests that the DS+ and CS gained comparable conditioned reinforcing properties.

## Experiment 2: Discriminative *vs* conditioned stimulus effects on sucrose-seeking

### IntA sucrose self-administration with discriminative stimuli

Figs. 4C-D show sucrose self-administration behaviour across the 20 IntA sessions. During Sessions 1-10, DS-periods were of fixed duration (i.e., 25 min), and during sessions 11-20 DS-periods lasted 20, 25 or 30 min. Across sessions, rats showed higher rates of responding on the active lever during DS+ periods than during DS-periods and responding also varied across sessions (Fig. 4C; DS type; *F* (1, 6) = 181.1, *p* < 0.0001; Session x DS type; *F* (19, 114) = 3.68, *p* < 0.0001). The rats did not escalate their sucrose intake across the 20 IntA sessions (Fig. 4D; Session effect; *F* (19, 120) = 1.41, *p* = 0.13). Thus, rats learned to respond for sucrose during DS+ periods and to suppress their responding during DS-periods.

Figures 4E-H show the progression of this discrimination training across sessions by illustrating the rate of responding on the levers during DS+ and DS-presentations. On Session 1, the rats pressed more on the active vs. inactive lever (Fig. 4E; Lever type; *F* (1, 6) = 42.47, *p* = 0.0006), and the rate of lever pressing varied across the session (Time effect; *F* (234, 1434) = 3.85, *p* < 0.0001). On Session 1, rats were already making the majority of their active lever presses during DS+ presentations (pie chart inset in Fig. 4E; 80 % for DS+ and 20 % for DS-, SEM ± 1.17; *t* (6) = 25.54, *p* < 0.0001). This pattern of discriminated responding persisted through Session 10 (Fig. 4F; Lever type; *F* (240, 3120) = 4.5, *p* < 0.0001; Time effect; *F* (239, 1434) = 3.85 *p* < 0.0001; pie chart inset; 97 % for DS+ and 3 % for DS-, SEM ± 0.46; *t* (6) = 102.5 *p* < 0.0001).

As seen in Exp. 1 with cocaine, responding peaked during DS+ presentation and dipped during DS-presentation even when we varied the duration of the DS-periods (sessions 11-20; Figs. 4G-H). On the first such session (Session 11), responding was still highest during DS+ periods, with rats almost completely suppressing their active lever pressing during DS-periods (Fig. 4G; Lever type; *F* (1, 6) = 71.98, *p* = 0.0001; Time effect, *F* (239, 1434) = 7.21, *p* < 0.0001; pie chart inset; 99 % for DS+ and 1 % for DS-, SEM

± 0.23; *t* (6) = 208.5, *p* < 0.0001). Rats maintained this pattern of discriminated responding through Session 20 (Fig. 4H; Lever type; *F* (1, 6) = 118.1, *p* < 0.0001; Time effect, *F* (239, 1434) = 10.55, *p* < 0.0001; pie chart inset; 99 % for DS+ and 1 % for DS-, SEM ± 0.35; *t* (6) = 138.8, *p* < 0.0001). Thus, during IntA sessions, rats were not pacing their sucrose self-administration behaviour using an internal timer. Instead, sucrose self-administration behaviour was under the control of the discriminative stimuli signaling when sucrose was available vs. unavailable.

### Cue-induced sucrose seeking

Two and 21 days following the last IntA session (session 20), we assessed sucrose-seeking behaviour induced by response independent presentation of the DS+, DS-, CS and DS+CS combined. Each cue was presented 4 times on each test session, and lever pressing produced no outcome (Fig. 5A; no sucrose or cues). We analysed the rate of responding to the first presentation of each cue (Figs. 5B and D) and across the 4 presentations of each cue (Figs. 5C and E).

On Day 2 of forced abstinence, rats responded most on the previously sucrose-associated (active) lever, and this effect depended on cue type (Fig. 5B shows active lever presses and Supplemental Fig. 2A shows inactive lever presses; Lever type, *F* (1, 6) = 6.96, *p* = 0.04; Cue type *F* (4, 24) = 4.49, *p* = 0.007; Lever x Cue *F* (4, 24) = 4.26, *p* = 0.01); Fig. 5C shows active lever presses and Supplemental Fig. 2B shows inactive lever presses; Lever type, *F* (1, 6) = 6.64, *p* = 0.04; Cue type *F* (4, 24) = 5.61, *p* = 0.002; Lever x Cue effect *F* (4, 24) = 4.88, *p* = 0.005). Cue type did not influence inactive lever presses (Supplemental Figs. 2A-B; all *p* values > 0.99). However, cue type determined responding on the previously sucrose-associated lever. First, only DS+ and DS+CS presentation increased responding on the previously sucrose-associated lever compared to responding during the inter-trial interval (Fig 5B; DS+ v. ITI, *p* = 0.0008, Fig 5C; DS+ v. ITI). Second, DS+ and DS+CS presentation also produced the highest rates of responding compared to other cue types (Fig 5B; DS+ v. CS, *p* = 0.0008; DS+ v. DS-, *p* = 0.0008; Fig 5C; DS+ v. DS-, *p* = 0.0008; DS+ v. CS, *p* = 0.0008; DS+CS v. DS-, *p* = 0.006; DS+CS v. CS, *p* = 0.006).

On Day 21 of forced abstinence, analysis of the response to the first presentation of each cue showed no significant main or interaction effects (Fig. 5D; Lever type, *F* (1, 6) = 4.48, *p* = 0.08; Cue effect *F* (4. 24) = 1.62, *p* = 0.2; Lever x Cue effect, *F* (4, 24) = 2.21, *p* = 0.1). However, analysis of responding across all 4 presentations of each cue type showed that rats responded most on the previously sucrose-associated (active) lever, and this effect depended on cue type (Fig. 5E; Lever type, *F* (1, 6) = 9.87, *p* = 0.02; Cue effect *F* (4. 24) = 2.40, *p* = 0.08; Lever x Cue effect, *F* (4, 24) = 2.89, *p* = 0.04). Further analysis of this interaction effect showed that DS+ presentation produced greater rates of responding on the previously sucrose-associated lever as compared to the DS- or CS (Fig. 5E; DS+ v. DS-; *p* = 0.03; DS+ v. CS, *p* = 0.02). Cue type did not influence inactive lever presses (Supplemental Figs. 2C-D; all *p* values > 0.99).

## Experiment 3: Conditioned inhibition of cocaine seeking

### IntA cocaine self-administration with discriminative stimuli

Figs. 6C-D show cocaine self-administration behaviour across the 15 IntA sessions. Rats pressed more on the active lever during DS+ vs. DS-periods, and this discrimination effect increased across sessions (Fig. 6C; DS type; *F* (1, 40) = 23.3, *p* < 0.0001; Session x DS type; *F* (14, 560) = 3.86, *p* < 0.0001). Cocaine intake also escalated across the 15 sessions (Fig. 6D; Session effect; F (14, 280) = 9.09, *p* < 0.0001; *t* (20) = 5.92; Session 1 vs. Session 15; *t* (20) = 5.92, *p* < 0.0001), increasing by 905% (SEM ± 188) from Session 1 to 15. Thus, rats progressively learned to respond for cocaine during the DS+ and to inhibit their responding during the DS-.

Figs. 6E-F illustrate the rate of responding on the levers on the first v. last IntA session as a function of DS+ and DS-presentation. On Session 1, the rats pressed similarly on the active vs. inactive lever (Fig. 6E; Lever type; *F* (1, 20) = 0.62, *p* = 0.44) and responding varied across the session (Time effect; *F* (239, 4780) = 2.24, *p* < 0.0001). Active lever presses, which were low on Session 1, were also relatively evenly distributed across DS+ vs. DS-periods (pie chart inset in Fig. 6E; 55 % for DS+ and 45 % for DS-, SEM ± 5.97; *t* (20) = 0.80, *p* = 0.43). By Session 15, rats increased their responding and pressed significantly more on the active vs. inactive lever, and lever pressing also varied across the session (Fig. 6F; Lever type; *F* (1, 20) = 22.25, *p* = 0.0001; Time effect; F (239, 4780) = 8.06, *p* < 0.0001). Notably, by Session 15, rats made most of their active lever presses during DS+ periods (pie chart inset in Fig. 6F; 79 % for DS+ and 21 % for DS-, SEM ± 3.29; *t* (20) = 0.80 *p* < 0.0001). Thus, cocaine self-administration behaviour was under the control of discriminative stimuli signaling when the drug was available versus unavailable.

### Conditioned inhibition of cue-induced cocaine seeking

On Day 2 following the last IntA session (Session 15), we assessed cocaine-seeking behaviour induced by response-independent presentation of the DS+, DS-, CS and DS+DS-combined. Each cue was presented 4 times on each test session, and lever pressing produced no outcome (Fig. 7A; no cocaine or cues). We analysed the rate of responding to the first presentation of each cue (Fig. 7B) and across the 4 presentations of each cue (Fig. 7C). Rats responded most on the previously cocaine-associated (active) lever, and this effect depended on cue type (Fig. 7B shows active lever presses and Supplemental Fig. 3A shows inactive lever presses; Lever type, *F* (1, 20) = 46, *p* < 0.0001; Cue type *F* (4, 80) = 6.43, *p* = 0.0002; Lever x Cue *F* (4, 80) = 7.65, *p* < 0.0001; Fig. 7C shows active lever presses and Supplemental Fig. 3B shows inactive lever presses; Lever type, *F* (1, 20) = 43.66, *p* < 0.0001; Cue type *F* (4, 80) = 11.02, *p* < 0.0001; Lever x Cue effect *F* (4, 80) = 11.87, *p* < 0.0001). Cue type did not influence inactive lever presses (Supplemental Figs. 3A-B; all *p* values > 0.05). However, cue type determined responding on the previously cocaine-associated lever. First, both the DS+ and CS enhanced responding on the previously cocaine-associated lever above that seen during the inter-trial interval (Fig. 7B; DS+ v. ITI, *p* < 0.0001; Fig 7C; DS+ v. ITI, *p* < 0.0001; CS v. ITI, *p* = 0.004. Second, the DS+ triggered greater increases in cocaine-seeking compared to all other conditions (Fig. 7B; DS+ v. DS-, *p* < 0.0001; DS+ v. CS, *p* = 0.0003; Fig 7C; DS+ v. DS-, *p* < 0.0001; DS+ v. CS, *p* = 0.003. In Fig. 7C, the CS also triggered more active lever pressing compared to both DS-, *p* = 0.0004 and DS+DS-, *p* = 0.002). Importantly, presenting the DS-together with the DS+ inhibited cocaine-seeking otherwise triggered by the DS+ (Fig. 7B; DS+DS-v. DS+, *p* < 0.0001; Fig 7C; DS+DS-v.DS+, *p* < 0.0001). Thus, presenting a DS-along with a DS+ prevents the DS+ from sparking cocaine-seeking behaviour.

## Discussion

Both conditioned stimuli (CSs) signaling reward delivery and discriminative stimuli (DSs) signaling reward availability can influence drug craving and relapse after abstinence. However, DSs are the only class of drug-associated cues present before and during drug relapse. Yet, the mechanisms of action of DSs are largely unknown. Here we compared CS and DS effects on cocaine-as well as sucrose-seeking behaviour following a history of intermittent-access self-administration, which is thought to model the intermittent patterns of cocaine use in humans. We report three main findings. Firstly, in female and male rats, discriminative stimuli (a DS+ and a DS-) came to tightly control cocaine (Exps. 1 & 3) and sucrose (Exp. 2) self-administration behaviour during intermittent-access sessions. Secondly, on days 2 and 21 of abstinence, the DS+ triggered increased cocaine- or sucrose-seeking behaviour, while the CS was ineffective. Thirdly, after 21 days of abstinence, the DS+ and CS, presented alone or together, equally reinforced instrumental responding under extinction (no cocaine). Finally, in Exp. 3, the DS-inhibited increases in cocaine-seeking behaviour otherwise triggered by the DS+, indicating conditioned inhibition of cocaine seeking. Thus, after intermittent-access self-administration, both DSs and CSs acquire similar conditioned reinforcing properties, but DSs surpass CSs in triggering reward-seeking behaviour.

### Discriminative stimulus control of IntA cocaine and sucrose self-administration behaviour

During intermittent-access sessions, rats had continuous access to two levers, and pressing the active lever delivered cocaine or sucrose during DS+ periods, but not during DS-periods. The rats developed a discriminative pattern of responding, pressing the active lever most for cocaine (Exps. 1 & 3) and sucrose (Exp. 2) during the DS+ and suppressing responding during the DS-. Indeed, across experiments, rats were making 79 to 99% of their total active lever presses during DS+ presentation. Our IntA discrimination procedure involves reinforced DS+ periods immediately before each nonreinforced DS-period. As such, cocaine concentrations would be high immediately before the start of each DS-period and start to dip during the DS-period [for cocaine modeling during IntA self-administration sessions see (Algallal et al. 2020; Allain et al. 2017a; Allain et al. 2017b; Kawa et al. 2019a)]. Prior work shows that cocaine self-administration increases when drug concentrations start to dip below an animal’s preferred level (Ahmed and Koob 2005; Zimmer et al. 2011). This suggests that during the DS+, rats learned to load up on cocaine and during the DS-, they learned to inhibit cocaine seeking, when it would normally be particularly pronounced. This aligns with past research showing discriminative stimulus control of drug self-administration behaviour (Ciccocioppo et al. 2001; Falk 1994; Falk and Lau 1995; McFarland and Ettenberg 1997; Weiss et al. 2000), including recent work using intermittent access procedures (Collins et al. 2022; Martin et al. 2022; Pitchers et al. 2017). However, in these recent IntA studies (Collins et al. 2022; Martin et al. 2022; Pitchers et al. 2017), the two DS types were presented in predictable sequence. As such, the apparent discriminated pattern of responding might involve rats tracking time. Our present results argue against a time-tracking explanation, because when we varied the length of DS-periods (20, 25 or 30 min) during IntA sessions, the rats consistently suppressed their responding during the DS-, regardless of period length. The rats immediately adapted to the change in DS length, also making an argument against habit-based cocaine (or sucrose) seeking (Everitt 2014). Thus, our findings suggest that through associative learning, discriminative stimuli control self-administration behaviour under intermittent-access conditions.

### Effects of conditioned versus discriminative stimuli on reward-seeking behaviour

At early and late abstinence times, we assessed cue-triggered increases in reward seeking. To this end, we exposed rats to cocaine- or sucrose-associated CSs and DSs independently of their behaviour, and we measured lever pressing in response to the cues. On both abstinence days 2 and 21, presentation of the DS+ alone or together with the CS significantly increased lever pressing. However, presenting the CS alone was ineffective. This suggests that through associative learning, cues indicating reward availability (DS+) may develop stronger incentive motivational properties than cues signaling reward delivery (CS), allowing drug-associated DSs to elicit strong craving and relapse (Robinson and Berridge 1993). This is relevant to humans, where drug cravings are most intense when discriminative stimuli signal imminent drug availability (Dar et al. 2010).

Some studies show that the ability of DSs to increase drug seeking persists or even incubates with prolonged abstinence and repeated testing (Ciccocioppo et al. 2004; Madangopal et al. 2019; Martin-Fardon and Weiss 2017; Weiss et al. 2001). Other research suggests that this effect diminishes with time and repeated testing, partially extinguishing during the first test session (Alleweireldt et al. 2001). In our experiments, cocaine (and sucrose) seeking triggered by the DS+ did not incubate with time; if anything, it started to diminish (e.g., compare rates of responding to the DS+ in Figs. 2B and 2D). Our procedures differ from previous studies in several ways. Some prior studies used trial-based discrimination training, whereby rats were exposed to 30 DS+ presentations and 30 DS-presentations on each session for example (Madangopal et al. 2019). This is considerably more than the 8 presentations of each DS type we used. We also used IntA self-administration. Previous studies showing persistent or incubating DS+ effects involved self-administration sessions with more continuous drug access. However, IntA procedures do not preclude incubation of drug craving; we (Gueye et al. 2019) and others (Nicolas et al. 2019) have shown incubation of CS-induced cocaine seeking after IntA cocaine use. Future work can assess the experimental parameters that determine how DS+ effects change with repeated testing and prolonged abstinence.

Our findings align with studies showing that passive exposure to a drug-associated DS increases drug seeking (Alleweireldt et al. 2001; Weiss et al. 2001), while passive exposure to a cocaine-paired CS is ineffective (Di Ciano and Everitt 2003; Grimm et al. 2000; Meil and See 1996; Weissenborn et al. 1995). However, these studies involved extinction training before drug-seeking tests. The relevance of extinction to humans is questionable (Epstein and Preston 2003; Fuchs et al. 2008), and the presence or absence of extinction training can also make a difference in terms of underlying neural substrates (Augur et al. 2016; Fuchs et al. 2006; Muller Ewald et al. 2019). Our procedure offers a way to compare the effects of response-independent exposure to CSs and DSs on relapse behaviour, without extinction training. In addition to the absence of extinction training, our procedure also involves a clinically relevant intermittent drug self-administration paradigm [(see also (Martin et al. 2022)]. Developing animal models that more closely mimic real-world scenarios is crucial for better understanding the neurobiological, psychological, and behavioral processes through which exposure to drug-associated cues triggers cravings and drug use in addiction.

### Conditioned inhibition of cocaine seeking

Environmental cues indicating the absence of drugs can work as conditioned inhibitors of drug seeking (Kearns et al. 2005; Madangopal et al. 2021; Mihindou et al. 2013; Rescorla 1969). Our work builds on existing literature by exploring this using an IntA cocaine self-administration procedure to better replicate the intermittency of cocaine use in humans. After IntA cocaine experience, we found that presenting a cue signaling drug absence (DS-) alongside a cue signaling drug availability (DS+) reduced the excitatory effects of the DS+ on drug seeking. Developing strategies to enhance inhibitory control over cocaine use has clinical significance. For example, integrating conditioned inhibitors of drug seeking, like an object or thought, into cue-exposure/regulation therapy could enhance outcomes by giving people with addiction an active means to deal with excitatory drug cues and prevent relapse (Kearns et al. 2005).

### Conditioned and discriminative stimuli acquire similar conditioned reinforcing value

Drug users encounter drug cues that result from their drug seeking actions (i.e., response-dependent) or that are unrelated to those actions (i.e., response-independent), and both experiences contribute to the persistent vulnerability to relapse during periods of abstinence (Brandon et al. 2007). For example, the sensation of throat numbness after snorting cocaine is a drug cue that results from drug seeking, while running into a former drug-taking partner can be a cue unrelated to any drug seeking action. Both experiences can trigger cravings and drug use. In laboratory rats, the impact of a drug-paired CS on relapse is usually studied by presenting the CS contingent on a response, where rats can press a lever to produce the CS without drug after a period of abstinence (Davis and Smith 1976; de Wit and Stewart 1981; Gueye et al. 2019; Meil and See 1996; 1997; Shaham et al. 2003; Shalev et al. 2002). In these conditions, the CS maintains drug-seeking actions through conditioned reinforcement (Robinson and Berridge 1993). Here we found that after IntA cocaine exposure, a DS and a CS associated with drug use acquired comparable conditioned reinforcing properties. This allowed the cues to reinforce the lever-pressing response that previously produced cocaine. Therefore, following a history of intermittent cocaine use, drug-associated DSs and CSs can be equally effective in reinforcing and sustaining cocaine-seeking actions [see also (Di Ciano and Everitt 2003)]. In contrast, response-independent presentation of a cocaine- or sucrose-associated DS significantly increased reward-seeking behaviour, while a reward-associated CS did not. Thus, our findings support the idea that CSs and DSs influence reward-seeking actions through at least partially distinct learning and motivational mechanisms (Di Ciano and Everitt 2003).

## Conclusions

Conditioned stimuli associated with drugs and discriminative stimuli indicating drug availability both play a role in relapse. However, we know less about how discriminative stimuli work, even though they control drug seeking before and during relapse, serving as triggers for cravings and relapse. In line with recommendations to develop science-based treatments for addiction (Epstein et al. 2018), we have refined a behavioural paradigm to distinguish between the effects of discriminative and conditioned stimuli in the same individuals, following a history of intermittent cocaine intake, which reflects the intermittency of human cocaine use (Allain et al. 2015; Beveridge et al. 2012; Cohen 1994; Leri et al. 2004). Our results show that both cue types can acquire similar conditioned reinforcing value. However, discriminative stimuli are notably more effective in sparking the desire for cocaine or sucrose reward. Thus, cues predicting reward availability may engage unique motivational and neurobiological processes to influence reward-seeking behaviour.

## Acknowledgements

We thank Drs. Terry E. Robinson, Donna J. Calu and David A. Martin for advice on our experimental protocol.

## Author Contributions

NAN, MJFR and ANS designed the research. NAN performed all experiments with assistance from SAS, DC, SCO, and IL. NAN analysed all data with guidance from ANS and assistance from SAS and MJFR. NAN and ANS wrote the article with revisions from co-authors.

## Note

The authors declare using ChatGPT to improve writing clarity in the Abstract text.

---

**Supplemental Fig. 1:**
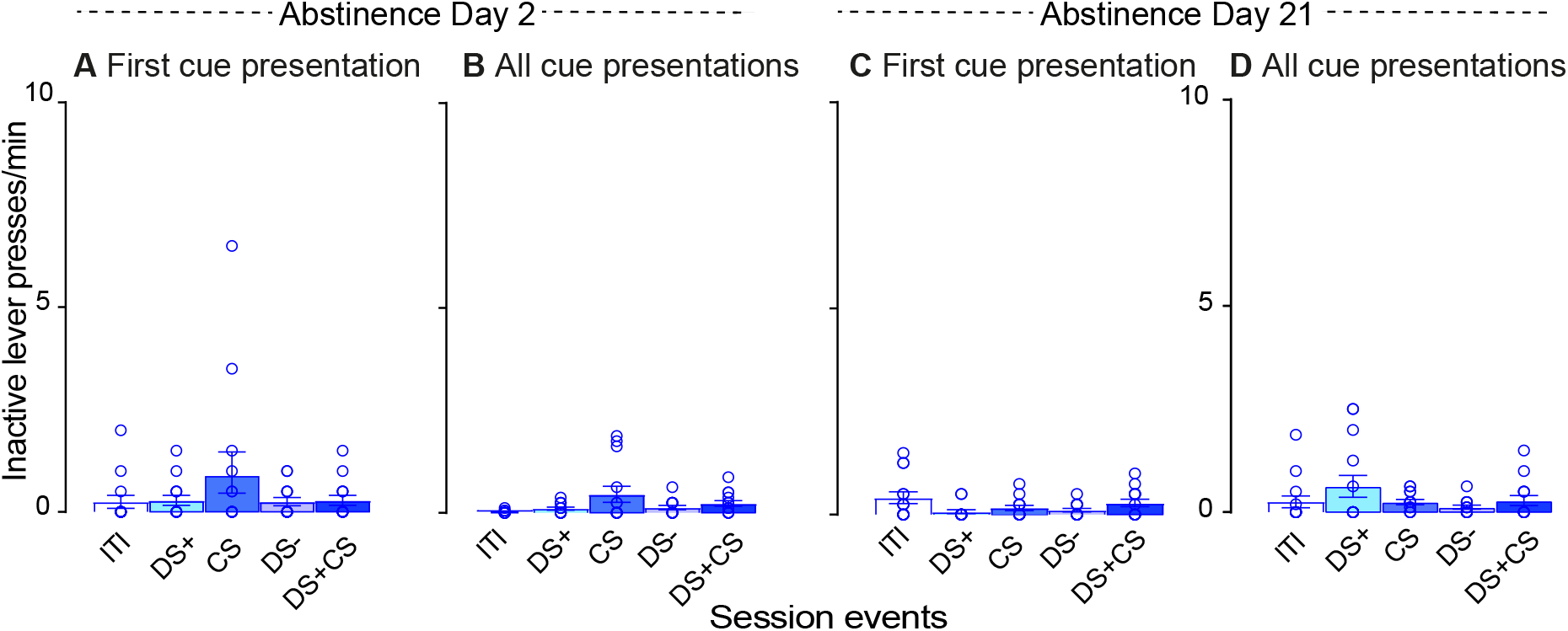
In Experiment 1, cue type did not influence responding on the inactive lever during cue-induced cocaine-seeking tests. **A** Inactive lever presses in response to the first presentation of each cue type. **B** Inactive lever presses across all 4 presentations of each cue type. ITI; inter-trial interval, DS; discriminative stimulus; CS; conditioned stimulus. Data are mean ± SEM, n = 14.

**Supplemental Fig. 2:**
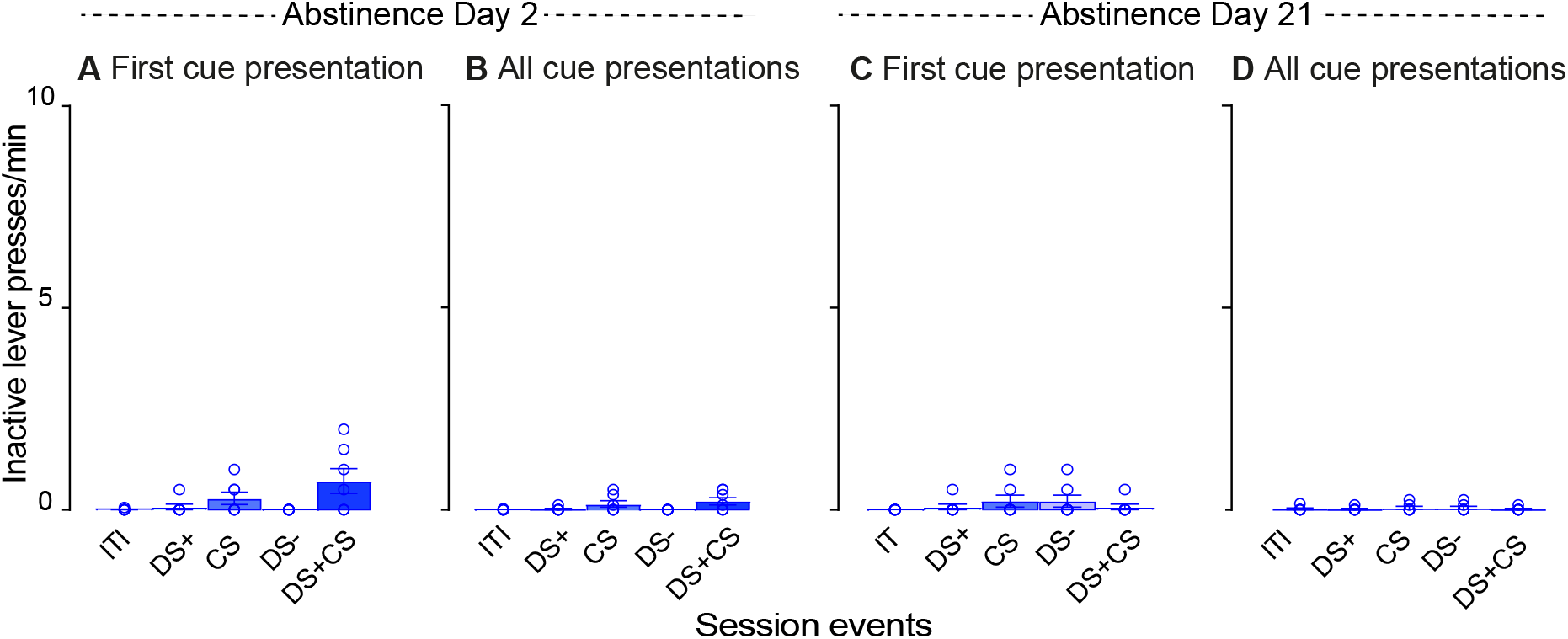
In Experiment 2, cue type did not influence responding on the inactive lever during cue-induced sucrose-seeking tests. **A** Inactive lever presses in response to the first presentation of each cue type. **B** Inactive lever presses across all 4 presentations of each cue type. ITI; inter-trial interval, DS; discriminative stimulus; CS; conditioned stimulus. Data are mean ± SEM, n = 7.

**Supplemental Fig. 3:**
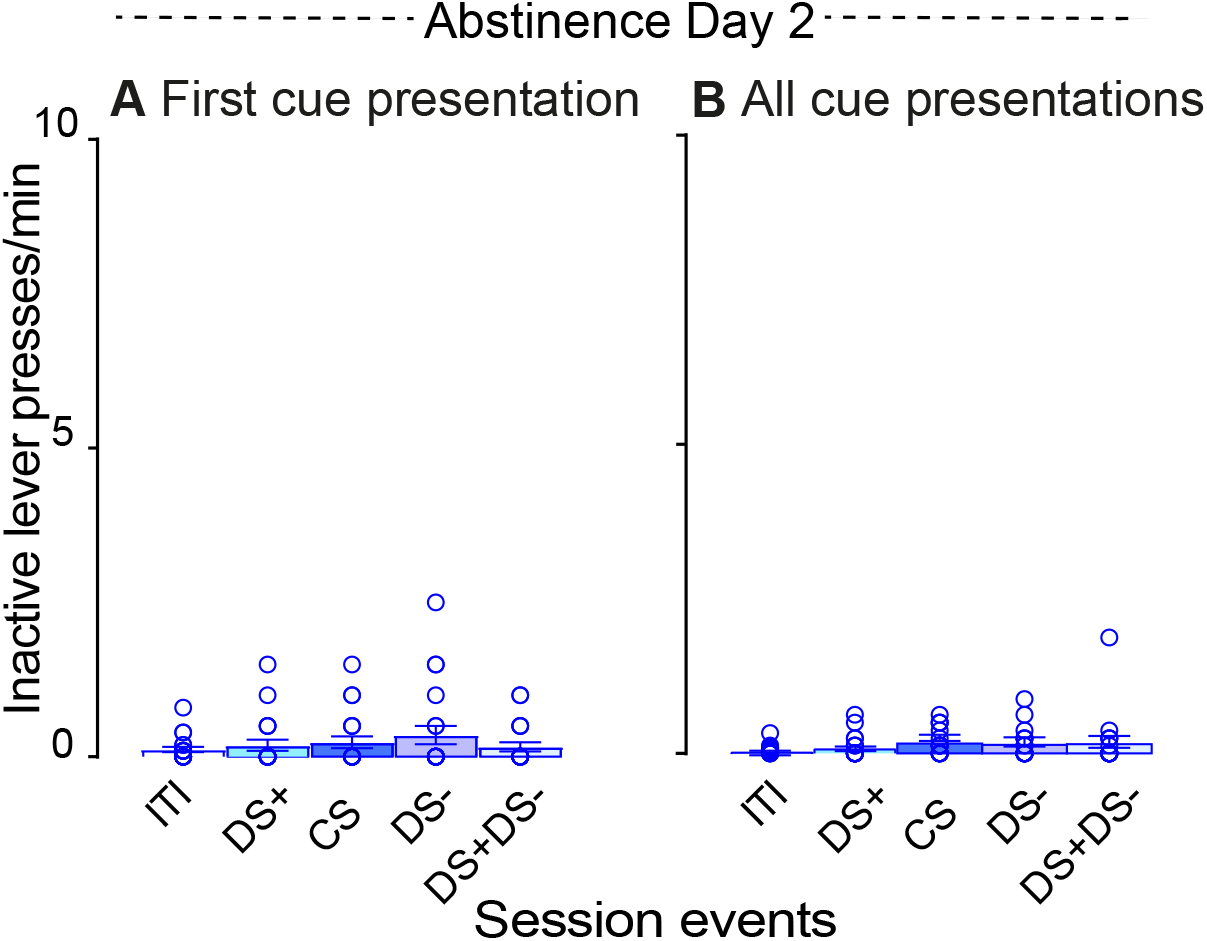
In Experiment 3, cue type did not influence responding on the inactive lever during cue-induced cocaine-seeking tests. **A** Inactive lever presses in response to the first presentation of each cue type. **B** Inactive lever presses across all 4 presentations of each cue type. ITI; inter-trial interval, DS; discriminative stimulus; CS; conditioned stimulus. Data are mean ± SEM, n = 21.

